# Successful introduction of the Colour Test into inexperienced settings

**DOI:** 10.1101/2020.08.07.241158

**Authors:** Kadri Klaos, Anna Agejeva, Tiina Kummik, Sirje Laks, Olesja Remets, Sirje Sasi, Anneli Tann, Piret Viiklepp, Alan Altraja

## Abstract

Implementation of non-commercial in-house methods into routine clinical diagnostics becomes more challenging, because these methods are not internationally standardized, most of the research in that field is underfunded and recommendations for their use is lacking. We conducted a study, where all the technicians were previously unfamiliar to the Colour Test (CT), a colorimetric redox indicator and thin-layer agar based *Mycobacterium tuberculosis* complex diagnosis and direct drug susceptibility testing (DST) method, and tested whether the performance of this in-house method is dependent on experience of the laboratory personnel.

After a two-day hands-on training, six panels of 150 *M. tuberculosis* isolates were cultured onto CT plates prepared in-house by six technicians in two laboratories. Finally, triplicate readings of 900 CT plates resulted 18 DST patterns for each of the initial isolates. The results were compared to each other and the gold standard of BACTEC MGIT 960 DST results.

The median time to produce *M. tuberculosis* CT DST results for three antituberculosis drugs was 13 days. The overall ability to correctly define phenotypic resistance ranged from 94.7% for levofloxacin to 95.8% and 97.3% for isoniazid and rifampicin, respectively. The test specificities were even higher exceeding 97% for all three drugs tested. Interobserver agreement reached 100% in one of the laboratories and exceeded 97% for levofloxacin and 99% for isoniazid and rifampicin in the second laboratory.

The implementation of the CT into a new laboratory was straightforward with only minimal guidance. This study proves that the CT is highly reproducible and easily interpreted by previously inexperienced personnel.

## Introduction

*Mycobacterium tuberculosis* infection is one of the oldest and deadliest epidemics experienced by the mankind, currently causing over 1.2 million deaths annually [1, 2]. In 2015, the World Health Organization (WHO) released the End TB Strategy to eliminate tuberculosis (TB) during the following 15 years [3]. Despite a small annual reduction of new TB cases and recent release of new anti-TB drugs, the global milestone of reducing the number of TB-associated deaths by 35% compared to 2015 cannot be reached in most parts of the world by the end of 2020 [4].

Timely and dependable diagnosis of TB with drug susceptibility testing (DST) results is essential for a successful treatment outcome [5]. However, TB is still underdiagnosed in all high-TB burden countries, where 87% of the disease is estimated to be located [1]. The implementation of innovative molecular techniques like the GeneXpert MTB/RIF have had a tremendous impact on TB detection, but has also shown that these methods cannot be used in every setting due to high infrastructural requirements and significant financial ramifications [6–10]. Conventional TB diagnosis with the initiation of a correct treatment requires phenotypic drug susceptibility testing (DST) results [11, 12]. The current gold standard for DST *i.e.* the BACTEC Mycobacterium Growth Indicator Tube (MGIT) 960 system is a fast and sensitive method [13]. The downside of the MGIT 960 system is that it needs a pure culture as a basis of DST testing, and it can take months until a result is available [14]. In those parts of the world, where the MGIT 960 system is not available due to high cost or lack of infrastructure, the proportion method on Löwenstein-Jensen (LJ) media is used [15]. LJ media for DST is prepared in-house and the proportion method does not require specialized equipment other than a thermostat, but it takes even longer than the MGIT 960 system to produce the final DST results.

A few methods that can simultaneously detect bacterial growth and perform DST have been developed [16]. One of these is the microscopic observation drug susceptibility (MODS) method, which has been shown to be highly sensitive and specific in the diagnosis of multidrug resistant (MDR)-TB from patient specimens [17, 18]. The only disadvantage of MODS is that it uses liquid medium in a 6-well plate that is put into a plastic bag. The plate is removed from the bag in order to read the results and therefore, the method is more prone to accidents than is a closed MGIT tube.

The thin-layer agar Colour Test (CT) is one of the in-house methods for DST that uses solid media [16]. The principle of this method lays on early microcolony detection and the proportion method for DST interpretation. CT combines thin-layer agar and colorimetric redox indicator-based principles as the agar media includes a redox indicator 2,-3-diphenyl-5-(2-thienyl) tetrazolium chloride that colours the microcolonies red and makes them visible to the naked eye when approximately 1 mm in diameter [19–21]. Bacteria of the *M. tuberculosis* complex (MTBC) have a distinct colony morphology on agar media and therefore, no further identification tests are needed [22]. CT consists of a four-quadrant Petri dish of which one quadrant serves as a growth control with no anti-TB drugs and three quadrants are supplemented with food colouring and used for DST to several drugs, e.g. for isoniazid (INH), rifampicin (RIF) and ciprofloxacin (CFX) [23]. In former studies, where the performance of CT was evaluated, the media was prepared in an expert laboratory elsewhere and transported into the study site or the specimens from the study site were transported into an expert laboratory for testing [24, 25]. Hence, the preparations of the media represent a constraint of the CT method. On the other hand, the performance of CT has been evaluated in the role of a direct test for both detection of TB and DST [25–28]. Furthermore, CT has been studied as a method for indirect DST on multidrug resistant (MDR)-TB isolates with promising results [23, 29]. Apart from that, CT has not been evaluated in inexperienced settings coupled with on-site preparation of the media.

To fill this gap, we aimed at assessing the reproducibility, objectivity, accuracy and ease of implementing CT in DST of *M. tuberculosis* isolates in previously inexperienced settings, as one of the hurdles in implementing this technique might be the in-house preparations of the media.

## Materials and methods

### Ethics approval

Ethics approval for the study protocol was granted by the Research Ethics Committee of the University of Tartu, Tartu, Estonia with the protocol number 269/T-12. Consent was not sought from patients because data was analysed anonymously and only pure *M. tuberculosis* cultures and no patient specimen where used in this study.

### Study design

This study was conducted as a blinded retrospective study in two clinical laboratories in Estonia. The study lasted 6 months in the laboratory A (Lab A) (from September 2017-February 2018) and 7 months in the laboratory B (Lab B) (from October 2019-April 2020). All study technicians were previously completely unfamiliar with preparation of the CT media and interpretation of the DST results.

*M. tuberculosis* isolates were randomly selected from the Estonian TB archive from 2014 backwards until 2009. The archived isolates had previously obtained DST results from the BACTEC MGIT 960 system (Becton Dickinson and Co., Franklin Lakes, NJ, USA) according to the WHO standards and manufacturer’s recommendations [30, 31] with the following critical concentrations: INH 0.1 μg/mL, RIF 1.0 μg/mL and levofloxacin (LVX) 1.0 μg/mL.

All frozen samples from the archive where first sub-cultured into MGIT media and checked for contamination, when flagged positive. The positive liquid culture was divided into four cryovials (1 mL of culture each) and kept at 37°C for two weeks allowing the bacteria to populate before renumbering and freezing as three separate blinded samples for inexperienced study technicians in the Lab A and one back-up sample. The same batch of the three blinded study samples was used in the Lab B (Fig 1).

**Fig 1.**
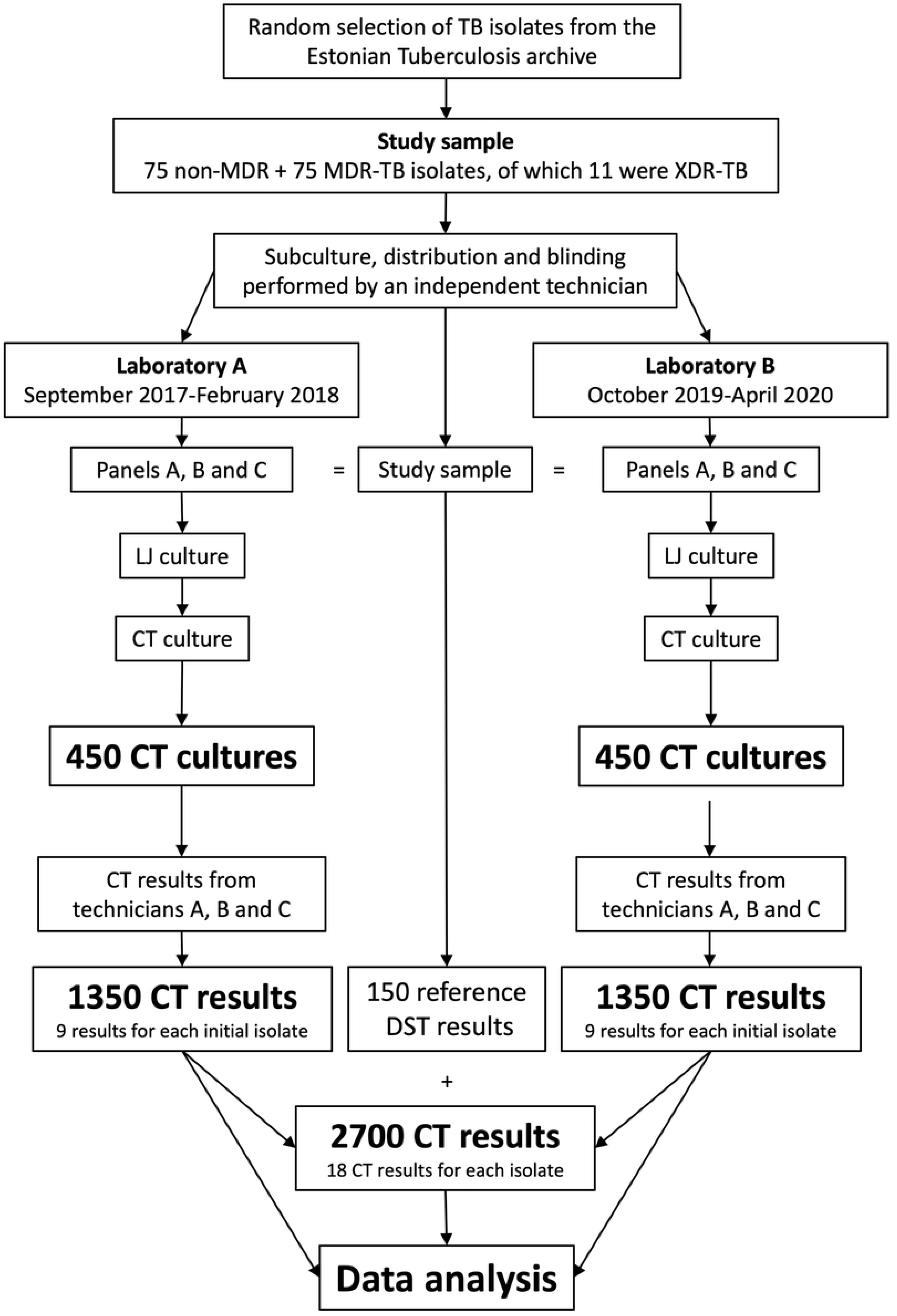
Flow chart of the study. Schematic view of the study process. TB - *Mycobacterium tuberculosis*; MDR - multidrug resistant; XDR - extensively drug resistant; LJ - Löwenstein-Jensen; CT - Colour Test; DST - drug susceptibility test.

Three technicians from both laboratories (six technicians in total) participated in this study. Prior to the study initiation, a two-day hands-on training was conducted at both study sites. After being acquainted to the CT media preparations, one of the laboratories decided to use a MediaJet machine (INTEGRA Biosciences AG, Zizers, Switzerland) to pour the plates, whereas the other laboratory continued to pour plates by hand with a Pipetboy (INTEGRA Biosciences AG, Zizers, Switzerland) and a 10 mL sterile serologic pipette. Workflow guidance was given to the technicians in both laboratories during first individual media preparations. Every technician first sub-cultured a portion (up to 1/3) of their samples onto LJ media and in the following week, he/she prepared four-quadrant Petri dishes with the CT media and performed sterility and growth controls. After two weeks, the LJ media grew positive with *M. tuberculosis* culture and the CT plates were inoculated with a suspension of the bacteria that were to be tested. Every technician also cross-read the CT plates cultured by his/her colleagues from the same laboratory. All results were photographed and the time to the result, colony counts, and DST results were documented for each plate.

The diagnostic performance of the CT was evaluated compared to the BACTEC MGIT 960 system DST results. All photographs of discordant results were re-evaluated, and the respective samples were retested with HAIN Genotype MTBDRplus (MTBDRplus) and MTBDRsl (MTBDRsl) (Hain Lifescience, Nehren, Germany) line-probe assays, when needed.

### Colour Test method

The CT plates were prepared by the individual technicians in the study laboratories as described previously [23] with the exception of using LVX (1 μg/mL) instead of ciprofloxacin as a proxy for fluoroquinolone resistance [11]. Each CT batch was quality-controlled using the *M. tuberculosis* H35Ra strain (ATCC 25177) and 3 sterility controls. Ready-to-use CT plates were stored at 4°C for 1 month.

In order to inoculate the CT plate, a strain suspension was made by inoculating a 1-μL inoculation loop full of fresh mycobacterial culture into a 5 mL tube with 0.5 mL of sterile water and glass beads. The tube was vortexed for 30 seconds and allowed to stand for at least 20 minutes allowing aerosols to descend. One mL of sterile water was added to the vortexed tube. The obtained suspension was further diluted to 1/5 in a new 5 mL tube. The CT plate was inoculated by adding one drop of the final suspension to each quadrant of the media and spreading with 10-μL loops. Every plate was sealed with parafilm and placed into a plastic zip-lock bag. All plates were incubated at 37°C for up to 6 weeks. The CT plates were read three times per week on Mondays, Wednesdays and Fridays. The DST results were interpreted, when at least 50 colonies appeared in the control quadrant. A strain was deemed resistant to a drug, when >1% of the colonies appeared in a drug quadrant, compared to the control quadrant. After 6 weeks or when the result was obtained and the plate was photographed, the plate was discarded.

### Statistical analysis

#### Calculation of the study sample

The optimal sample size was calculated according to Hajian-Tilaki taking into account a pre-determined sensitivity of at least 0.95, an α value of 0.05, a marginal distribution of not more than 0.05 and a prevalence of MDR of 0.5 according to the BACTEC MGIT 960 system [32].

#### Data analysis

All study results were entered into Microsoft Excel spreadsheets, unblinded and linked with MGIT 960 DST data obtained from the Estonian Tuberculosis Registry. The concordance of the CT DST results with those by the MGIT 960 system was assessed and the test performance characteristics were calculated. Intra- and inter-observer agreements were calculated as Fleiss’ Kappa [33].

The test performance characteristics (sensitivity, specificity, total agreement, and median time to result), as well as inter- and intra-observer agreements, were calculated using IBM SPSS Statistics software (version 20.0.0, Armonk, NY, USA). R: A language and environment for statistical computing (version 4.0.0, R Foundation for Statistical Computing, Vienna, Austria) with the “meta” package, was used to draw the forest plots [34].

## Results

Two lists were produced from archived MDR isolates, as well as the non-MDR isolates from 2009-2014 consisting of 413 and 1090 entries, respectively. Seventy-five entries from both lists were randomly selected resulting the study sample. The final study sample thus included 150 *M. tuberculosis* isolates. All of the selected archived TB isolates grew in MGIT media after initial thawing. Half of the 150 isolates selected were MDR and 11 (7.3%) of these were extensively drug resistant (XDR)-TB strains. Sixty-one isolates were pan-susceptible, 13 (8.7%) and 1 (0.7%) were INH- and RIF-monoresistant, respectively, according to the pre-analysis data from the Estonian Tuberculosis Registry obtained formerly with the BACTEC MGIT 960 system. Five (5.6%) of the 89 isolates with any resistance to anti-TB drugs did not have DST data on fluoroquinolone resistance and were therefore excluded from the respective analysis for the test performance.

A total of 20 batches of CT media were prepared for this study (10 batches in both laboratories). Each technician prepared at least 3 batches of media. The respective study technicians in both laboratories expressed that they needed to be focused during the media preparations. The median time to prepare and pour one batch of media without the time for autoclaving was 112.5 min in both laboratories with an IQR of 91.25-123.75 min and 106.25-130 min for Lab A and Lab B, respectively. Each batch consisted of a median of 59 CT plates (IQR 55-60 plates) and the median time to pour one CT plate was 117.5 sec (IQR 107.7-140 sec). No contamination was observed in any of the cultured plates and all the positive controls grew in the control quadrant and had no growth in the respective drug quadrants. No issues were discovered in the plate inoculation or plate reading steps.

A total of 943 CT plates (461 in Lab A and 482 in Lab B) were inoculated during this study. Each of the plates was read 3 times per week until positive or negative for up to 6 weeks. It took a median of 153 sec (IQR 143-181 sec) in Lab A and 154 sec (IQR 143-157 sec) in Lab B for one technician to continuously read and document end results of one CT plate.

The median time until the DST result was 13 days (IQR 11-15) for two sites together and 12 (IQR 10-15) and 13 (IQR 11-15) days for Lab A and Lab B, respectively. Four out of six technicians had a median time to result of 12 days. The time to result ranged from 5 to 45 days (Fig 2). The result pictures of CT plates that grew ≥35 days showed that five out of six plates in Lab A (in total 15 readings) and five plates in Lab B (9 readings in total) were overgrown and could have been read earlier (Fig 3).

**Fig 2.**
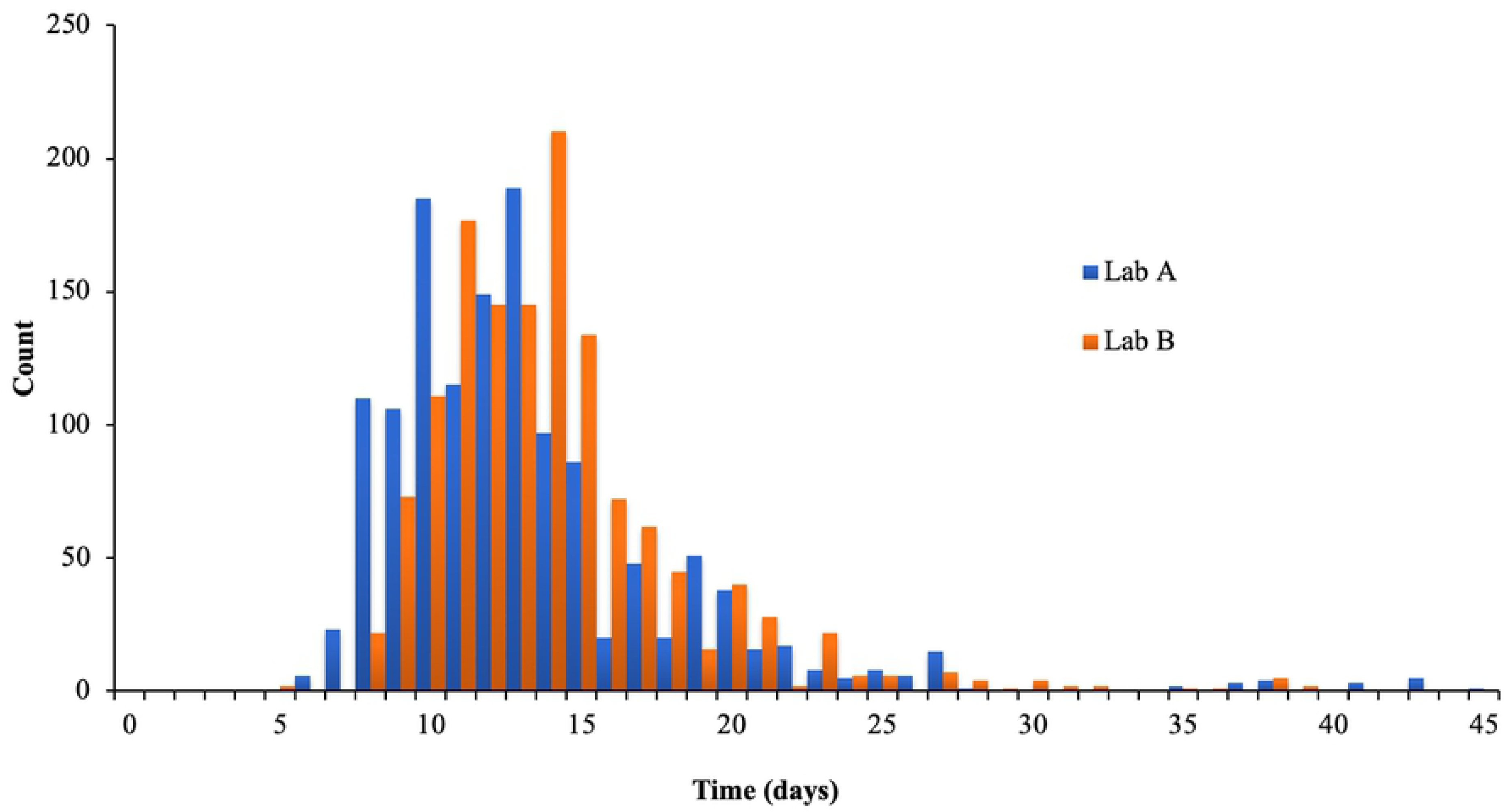
Time in days until a Colour Test result. A histogram was drawn from the times until a positive *Mycobacterium tuberculosis* Colour Test result in the laboratory A (Lab A) (blue, 1337 Colour Test readings) and the laboratory B (Lab B) (orange, 1347 Colour Test readings).

**Fig 3.**
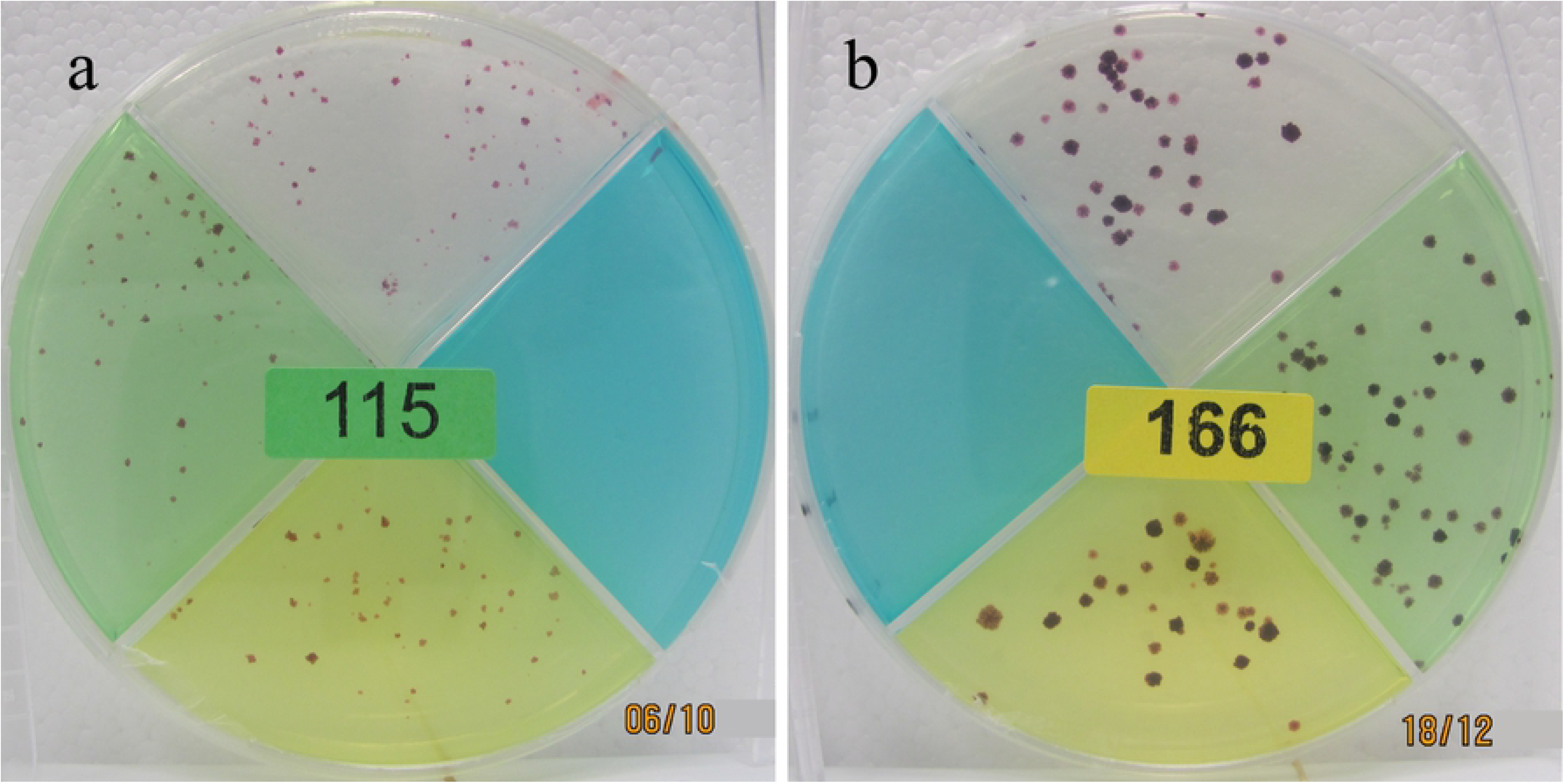
Photographs of positive Colour Test plates. Examples of *Mycobacterium tuberculosis* Colour Test results on day 14 (a) and 43 (b) after inoculation.

The results of five TB culture samples out of 900 (6 duplications of 150 samples) (0.6%) were excluded from the subsequent analysis due to insufficient growth on LJ media despite multiple attempts to culture (one isolate in Lab B) or in the CT control quadrant (four isolates from Lab A). For the following DST performance analysis, these 5 isolates were discarded and the CT DST results for 895 TB isolates were used for definite analyses.

The total agreement of the 2685 CT DST results with the reference method of MGIT 960 was 96.7, 97.2, and 97.8% for INH, RIF, and LVX, respectively (Table 1 and Fig 4). The pooled sensitivities for the detection of resistance to a certain drug ranged from 94.7% for LVX to 95.8% and 97.3% for INH and RIF, respectively. The specificity exceeded 97% for all drugs being 97.1, 98.1 and 98.2% for RIF, INH and LVX, respectively.

**Table 1.** Drug susceptibility testing performance of the *Mycobacterium tuberculosis* Colour Test compared to the MGIT 960.

|  |  |  | MGIT 960 |  | Total agreement<br>(95% CI) | Sensitivity<br>(95% CI) | Specificity<br>(95% CI) | LR+<br>(95% CI) |
| --- | --- | --- | --- | --- | --- | --- | --- | --- |
|  |  |  | S | R |  |  |  |  |
| CT<br>INH | Pooled | S | 1083 | 67 | 96.7%<br>(96.0-97.3) | 95.8%<br>(94.7-96.7) | 98.1%<br>(97.1-98.8) | 50.3<br>(33.0-76.9) |
|  |  | R | 21 | 1514 |  |  |  |  |
|  | Lab A | S | 537 | 42 | 96.0%<br>(94.8-97.0) | 94.7%<br>(92.9-96.1) | 97.8%<br>(96.2-98.9) | 43.3<br>(24.8-75.8) |
|  |  | R | 12 | 747 |  |  |  |  |
|  | Lab B | S | 546 | 25 | 97.5%<br>(96.5-98.3) | 96.8%<br>(95.4-98.0) | 98.4%<br>(96.9-99.3) | 59.7<br>(31.2-114.2) |
|  |  | R | 9 | 767 |  |  |  |  |
| CT<br>RIF | Pooled | S | 1281 | 37 | 97.2%<br>(96.5-97.8) | 97.3%<br>(96.3-98.1) | 97.1%<br>(96.0-97.9) | 32.9<br>(24.2-44.9) |
|  |  | R | 39 | 1328 |  |  |  |  |
|  | Lab A | S | 638 | 27 | 96.6%<br>(95.4-97.5) | 95.9%<br>(94.2-97.3) | 97.2%<br>(95.6-98.3) | 34.0<br>(21.8-53.0) |
|  |  | R | 19 | 654 |  |  |  |  |
|  | Lab B | S | 643 | 10 | 97.8%<br>(96.8-98.5) | 98.5%<br>(97.2-99.3) | 97.1%<br>(95.6-98.2) | 34.2<br>(22.2-52.6) |
|  |  | R | 20 | 674 |  |  |  |  |
| CT<br>LVX | Pooled | S | 2216 | 18 | 97.8%<br>(97.1-98.3) | 94.7%<br>(91.7-96.8) | 98.2%<br>(97.6-98.7) | 53.4<br>(39.2-72.7) |
|  |  | R | 40 | 321 |  |  |  |  |
|  | Lab A | S | 1104 | 0 | 98.4%<br>(97.5-99.0) | 100%<br>(97.8-100) | 98.1%<br>(97.2-98.8) | 53.6<br>(35.1-81.8) |
|  |  | R | 21 | 168 |  |  |  |  |
|  | Lab B | S | 1112 | 18 | 97.2%<br>(96.1-98.0) | 89.5%<br>(83.9-93.6) | 98.3%<br>(97.4-99.0) | 53.3<br>(34.0-83.4) |
|  |  | R | 19 | 153 |  |  |  |  |

Lab A and Lab B - laboratories; CT - Colour Test; MGIT - Mycobacterium Growth Indicator Tube; DST - drug susceptibility test; INH - isoniazid; RIF - rifampicin; LVX - levofloxacin; S - sensitive; R - resistant; CI - confidence interval; LR+ - Positive Likelihood Ratio; LR- - Negative Likelihood Ratio

**Fig 4.**
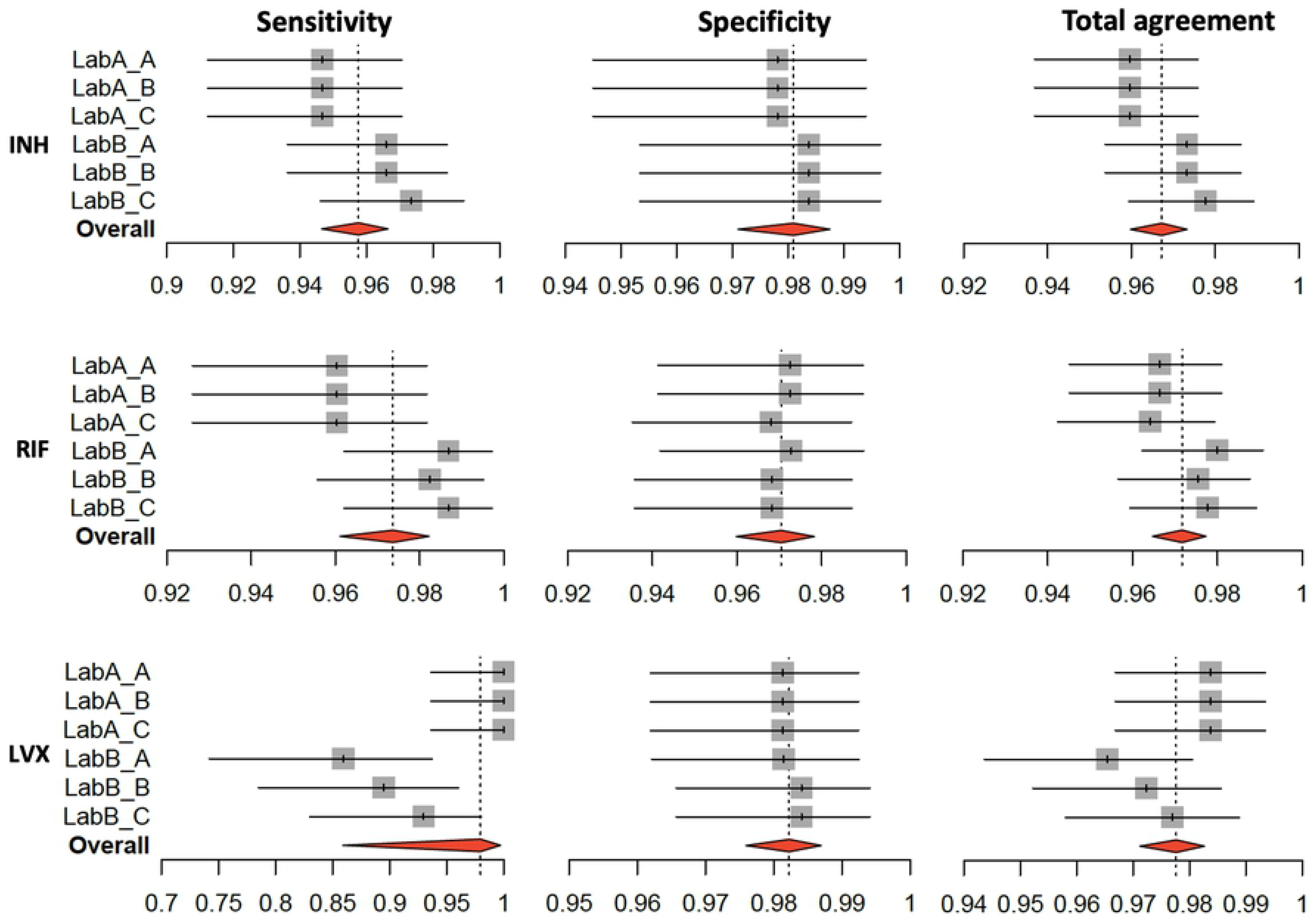
Individual and pooled sensitivities, specificities and total agreements of the Colour Test in DST of *Mycobacterium tuberculosis* isolates. Lab A - Laboratory A; Lab B - Laboratory B; INH - isoniazid; RIF - rifampicin; LVX - levofloxacin. Grey squares indicate the value for the respective technician; red diamonds indicate pooled estimates. During 18 different readings in two sites, a total of 20 isolates out of 150 produced at least one false positive or one false negative result, when compared to the gold standard of MGIT 960 system results. For seven isolates out of 20 a false negative DST result for LVX was recorded in Lab B. The result photographs for these isolates showed that on 6 CT plates, microcolonies were present in the LVX quadrant (Fig 5).

Seventy-seven percent of the discordant results were produced by eight initial isolates that all had discordant results for over 30% of the 18 readings (range 33.3-100%) that could not be resolved by re-evaluating the result photographs. Three of these isolates had discordant results in 100% of the readings. These eight isolates that produced discordant results were tested twice (once from the study culture and once from the initial archived culture) with the MTBDRplus and MTBDRsl line-probe assays (Table 2). The molecular DST results of three isolates (30, 48 and 82) that cumulatively produced 41.1% of the discordant results were in concordance with the CT results.

**Table 2.** Molecular drug susceptibility test results with HAIN MTBDRplus and MTBDRsl assays compared to the previous phenotypic results for those 8 *Mycobacterium tuberculosis* isolates, which currently produced discordant results between the Colour Test and the MGIT 960 system.

| Isolate No. | Estonian TB Registry data (MGIT 960) |  |  | HAIN MTBDRplus/sl |  |  | Colour Test |  |  |  |
| --- | --- | --- | --- | --- | --- | --- | --- | --- | --- | --- |
|  | INH | RIF | FLQ | INH | RIF | FLQ | INH | RIF | LVX | Count |
| 13 | R | R | S | R | S | S | R | R | S | 12 |
|  |  |  |  |  |  |  | S | S | S | 6 |
| 30 | R | S | S | R | R | R | R | R | R | 18 |
| 48 | S | S | S | R | R | R | R | R | R | 18 |
| 67 | R | R | S | S | S | S | S | S | S | 4 |
|  |  |  |  |  |  |  | S | R | S | 3 |
|  |  |  |  |  |  |  | R | R | S | 11 |
| 68 | R | R | S | S | S | S | S | S | S | 15 |
|  |  |  |  |  |  |  | R | R | S | 3 |
| 82 | R | S | S | S | S | NA | S | S | S | 18 |
| 117 | R | S | S | S | S | NA | S | S | S | 7 |
|  |  |  |  |  |  |  | R | S | S | 11 |
| 138 | R | R | S | S | S | S | S | S | S | 12 |
|  |  |  |  |  |  |  | R | R | S | 6 |

Count - the number of times a respective Colour Test result was observed out of 18 readings;
No. - isolate number (out of 150 samples); MGIT - Mycobacterium Growth Indicator Tube;
INH - isoniazid; RIF - rifampicin; FLQ - fluoroquinolone; S - sensitive; R - resistant; NA -
not applicable

The interobserver agreements between the three technicians in the Lab A, when reading DST results from 450 CT plates, was in 100% concordance (Table 3). In the Lab B, the Fleiss’ Kappa values for interobserver agreement were 0.99 for the DST results for INH and RIF and 0.97 for LVX. The intraobserver agreement, when reading results of triplicate testing of 150 isolates, was the same for all the technicians in Lab A ranging from 0.96 for INH to 0.97 for RIF and LVX DST results. The intraobserver agreement rates of the DST results in Lab B ranged from 0.85 for LVX to 0.97 for RIF. The intra-isolate agreements of the DST results for the 150 isolates including in total 18 readings of six CT plates were 0.95, 0.96 and 0.92 for INH, RIF and LVX, respectively.

**Table 3.** Fleiss’ Kappa values of inter- and intraobserver agreements. Interobserver agreement was calculated in each laboratory for 3 technicians reading 450 Colour Test plates. Intraobserver agreement was calculated for each technician reading triplicate Colour Tests of 150 isolates. Intra-isolate agreement was calculated for 18 readings of 6 duplications of Colour Tests for 150 isolates.

| Interobserver agreement<br>(450 CT plates; 3 technicians) |  |  |  |  |
| --- | --- | --- | --- | --- |
|  |  | Kappa | SD | P-value |
| Lab A | INH | 1.0 | 0.027 | <0.001 |
|  | RIF | 1.0 | 0.027 | <0.001 |
|  | LVX | 1.0 | 0.026 | <0.001 |
| Lab B | INH | 0.99 | 0.027 | <0.001 |
|  | RIF | 0.99 | 0.027 | <0.001 |
|  | LVX | 0.97 | 0.027 | <0.001 |
| Intraobserver agreement |  |  |  |  |

| **(150 isolates; 3 CT plates; 1 reader)** |  |  |  |  |
| --- | --- | --- | --- | --- |
| **Lab A**<br>**A, B, C** | INH | 0.96 | 0.046 | <0.001 |
|  | RIF | 0.97 | 0.046 | <0.001 |
|  | LVX | 0.97 | 0.045 | <0.001 |
| **Lab B**<br>**A** | INH | 0.95 | 0.047 | <0.001 |
|  | RIF | 0.97 | 0.047 | <0.001 |
|  | LVX | 0.85 | 0.046 | <0.001 |
| **Lab B**<br>**B** | INH | 0.95 | 0.047 | <0.001 |
|  | RIF | 0.96 | 0.047 | <0.001 |
|  | LVX | 0.89 | 0.047 | <0.001 |
| **Lab B**<br>**C** | INH | 0.96 | 0.047 | <0.001 |
|  | RIF | 0.96 | 0.047 | <0.001 |
|  | LVX | 0.91 | 0.047 | <0.001 |
| **Intra-isolate agreement**<br>**(150 isolates; 6 CT plates; 18 readings)** |  |  |  |  |
|  | INH | 0.95 | 0.006 | <0.001 |
|  | RIF | 0.96 | 0.006 | <0.001 |
|  | LVX | 0.92 | 0.006 | <0.001 |

CT - Colour Test; Lab A – Laboratory A; Lab B – Laboratory B; INH - isoniazid; RIF -
rifampicin; LVX - levofloxacin; SD - standard deviation

## Discussion

The commercial phenotypic DST methods for *M. tuberculosis* are well certified and provide reliable results but are not the best solutions for resource-constrained settings and laboratories with intense staff turnover. Instead, reliable and safer in-house methods could be used in order to allocate insufficient funding more effectively. The in-house methods including the CT come in handy, when a new economic crisis lies ahead, and supply chains work with long delays like they did in the European Union in spring 2020.

This is the first study to thoroughly evaluate the reproducibility, objectivity and ease of implementing the CT DST method into inexperienced settings with minimal guidance. To our knowledge, no other research has evaluated as many CT repetitions of one isolate as performed in this current study.

Two issues that were related to CT media preparations were discovered. First, the technicians needed to be focused during the four-quadrant Petri dish pouring phase in order to pour the drug-containing media with the correct sequence. The media was poured with a different sequence once during this study, but the problem was solved considering that different drug-containing media solutions are coloured with food colourings (i.e. INH-green; RIF-yellow and LVX-blue) and recognizable. The second concern was that the CT plates had a short realization time of only one month. During this study, the media was freshly prepared and used approximately a week after production, but when the CT would be used routinely in a diagnostic laboratory using direct patient specimen, it would be helpful to have a longer expiration date in order to pour more plates at once. Other study groups have used CT plates with expiry terms even up to four months [25], yet it needs to be investigated, whether such a prolongation changes the diagnostic performance of the test when the media is prepared on site.

None of the CT plates from 20 batches of media made by 6 different technicians was contaminated during this study, which indicates that the CT media is robust enough to be used in resource-constrained settings. Based on the experience from our current study, it is probable that the media could even be stored at room temperatures for a short period of time (1 week) and freshly made on a weekly basis, but this issue still needs to be investigated on site. It was surprising that even though one of the laboratories decided to use a MediaJet machine to pour the four-quadrant Petri dishes, whereas the other did the same by hand, the median time to prepare and pour one batch of CT media was not different between both laboratories (112.5 min) indicating that apart from the pouring, the most time-consuming parts of the media preparations lie in the phase, where the autoclaved and still warm Middlebrook 7H11 agar is supplemented by hand with antimycobacterial drug working solutions and food colourings.

This study showed that it took two and a half minutes in both laboratories to continuously read one CT plate until a result was documented. One could predict that this time could be slightly different and perhaps even shorter in routine settings, where the majority of cultured plates remain negative for six weeks and can quickly be examined three times weekly. The overall median time until a CT DST result was 13 days, which is consistent with previous results by others [23, 35]. Still, we found that a few of the CT plates were read very late, even 45 days after culturing (Fig 2 and 3) because our study protocol for indirect DST using the CT plates insisted of having 50 colonies in the control quadrant before reporting the results. In addition, the protocol for the CT with direct patient specimen does not assume the condition that the control quadrant needs to have at least 50 colonies before reporting DST and confirming MTBC and can therefore give the results faster.

The total agreement between the Colour Test and the MGIT 960 system was excellent with over 95% for all the anti-TB drugs tested (Fig 4). Furthermore, the specificities found in this study were > 97% for all technicians, which is in good correlation with previous studies using similar methods [23, 24, 28, 36]. The sensitivities varied between the study laboratories being slightly higher in the Lab B for the detection of INH and RIF resistance and higher in the Lab A with 100% for the detection of LVX resistance. The result photographs showed that in the Lab B microcolonies were present in LVX quadrants with false-negative results and the technicians did not take these as evidence of growth. The respective microcolonies did not have the cauliflower-like cording and were smaller than the colonies in other quadrants (Fig 5).

**Fig 5.**
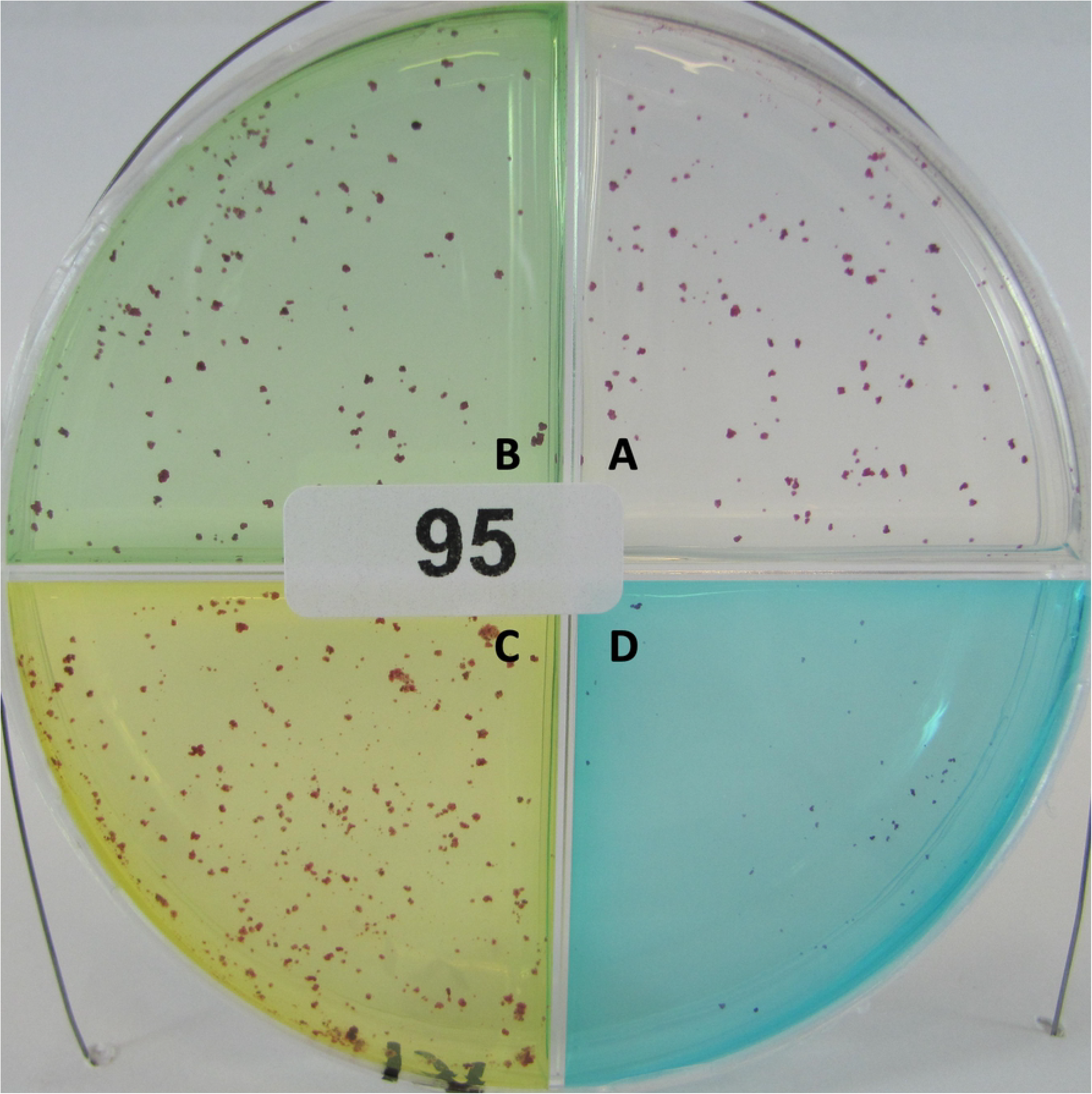
An example of a *Mycobacterium tuberculosis* Colour Test plate with microcolonies in the LVX quadrant. A - Growth control (colourless); B - INH quadrant (green); C - RIF quadrant (yellow); D - LVX quadrant (blue).

False negative or false positive DST results, when the MGIT 960 system was used as a reference, were recorded for 20 isolates. Three isolates out of 20 produced a discordant result 100% of times, i.e. on all 18 readings of 6 plates and MTBDRplus and/or MTBDRsl results were in concordance with the CT DST results for these isolates. These three isolates were the reason for over 40% of all discordant results. One explanation for the discordant result of the isolate 82 might be that the INH resistance was caused by a mutation in the *inhA* coding region that is not detected by the MTBDRplus but is detected by MGIT960 system with an INH critical concentration of 0.1 μg/mL (Table 2). Some mutations in the *inhA* are shown to cause low-level resistance to INH, and it might be possible that the critical concentration of 0.2 μg/mL in CT is the minimal inhibitory concentration for this strain [37]. The isolate number 30 was a MTBC contamination that probably occurred during the initial sputum decontamination phase in the laboratory and it included multiple probes from one day that were archived and were not marked as contaminations. We did not find any explanation for the discordant results of the isolate number 48. The only explanation is that there was a mix-up of strains during the initial archiving phase, because this patient with the strain from isolate 48 did not have any issues with their treatment and was already culture negative after two months of treatment. The isolates number 67 and 68 derived from one initial patient and the respective sputa were collected on the same day. MTBDRplus was conducted on the initial sputum that resulted in isolate number 67 and this test detected resistance conferring mutations for both INH and RIF. A possibility exists that this was a mixed infection and the initial MTBDRplus performed directly on the decontaminated specimen did detect a minority resistant strain, but further subculturing decreased proportion of the resistant strain in the population and therefore, the CT and later MTBDRplus and MTBDRsl tests had difficulties in the detection of resistance.

The interobserver agreement rates found were close to 1.0 except for LVX in the Lab B (Table 3 and Fig 4). The difference for the DST results for LVX in the Lab B might be because one of the technicians in the Lab B read CT plates up to one week later and also had better concordance with the gold standard regarding the DST for LVX (Fig 4). The intraobserver agreements and the intra-isolate agreements were mainly over 0.9 except for that in case of LVX in the Lab B, where microcolonies were not considered as growth in some of the CT plates.

During our study, the CT gave reliable DST results for three of the most important drugs used to treat TB [38], but whether this is true when using direct patient specimen or if it is possible to conduct further in-house indirect DST methods using colonies from the CT plate control quadrant needs to be clarified in future studies.

In conclusion, our study shows that the CT is easily implemented into new settings with minimal cost and no compromise in either antimycobacterial DST or time to result. We also show that the CT gives consistently same results on repetitive culturing of one isolate and is therefore highly reproducible and not dependent on the experience of laboratory personnel experience and moreover the readings of one CT plate by different inexperienced readers are the same.

## Acknowledgement

The authors are cordially thankful for the skilful technical assistance of Mrs. M. Noorkõiv.

